# Functional brain network topology across the menstrual cycle is sex hormone dependent and correlates with the individual well-being

**DOI:** 10.1101/2020.11.01.363937

**Authors:** Marianna Liparoti, Emahnuel Troisi Lopez, Laura Sarno, Rosaria Rucco, Roberta Minino, Matteo Pesoli, Giuseppe Perruolo, Pietro Formisano, Fabio Lucidi, Giuseppe Sorrentino, Pierpaolo Sorrentino

## Abstract

The menstrual cycle is known to influence the behaviour. The neuronal bases of this phenomenon are poorly understood. We hypothesized that hormones, might affect the large-scale organization of the brain functional networks and that, in turn, such changes might have behavioural correlates in terms of the affective state. To test our hypothesis, we took advantage of magnetoencephalography to investigate brain topology in early follicular, ovulatory and luteal phases, in twenty-four naturally-cycling women without signs of anxiety and/or depression. We show that in the alpha band the betweenness centrality (BC) of the right posterior cingulate gyrus (PCG) during the ovulatory phase is increased and the rise is predicted by the levels of estradiol. We also demonstrate that the increase in the BC is related to improved subjective well-being that, in turn, is correlated to the estradiol levels. The increased topological centrality of the PCG during the ovulatory phase could have implications in reproductive psychology.

## Introduction

The brain, over the course of a lifetime, undergoes continuous and dynamic changes on multiple time scales^1^, both structurally and functionally. These changes can be induced both by pathological processes^2–4^, as well as physiological and environmental factors including, as an example, dietary or sleeping habits^5,6^.

Hormonal modulation is also capable of inducing changes in both structure and function. In particular, sex hormones play a pivotal role in the modulation of behaviour. The patterns of sex hormones undergo physiological changes throughout life. As soon as the prenatal period, the hypothalamic-pituitary-gonadal axis, produces gender differentiation. Phoenix et al.^7^ hypothesized that testosterone acts on the brain, causing permanent changes which affect neurobehavioral development. Sexual differentiation of the brain is not limited to the prenatal development but extends throughout puberty. During puberty, the hormonal changes contribute to morphological variations of the cortical and subcortical regions^8–10^ involved in sensorimotor processing, such as the thalamus and the caudate, as well as areas involved in emotion and memory processes, such as the amygdala and the hippocampus. It has been proposed that sex hormones fluctuations during puberty might be responsible of gender-related differences in the brain functioning^11^. Finally, a possible role of sex hormones on brain activity has been invoked in aging, with lower estrogens negatively affecting cognitive functions and memory^12^.

Unlike puberty or menopause, which are unique and non-repeatable processes, the menstrual cycle is the only sex hormone-related phenomenon that repeats itself cyclically, with periodical, coordinated variations of multiple hormones including estradiol, progesterone, Follicular Stimulant Hormone (FSH) and Luteinizing Hormone (LH). Such variations can induce a number of physical (acne, breast pain, cramps, headaches), vegetative (sleep and eating disorders)^13,14^ and psychopathological changes (anxiety, depression, moodiness)^15^.

A large number of women suffer from sex hormone dependent depressive disorders, including postpartum depression, peri-menopausal depression and premenstrual dysphoric disorders (PMDD)^16,17^. PMDD is characterized by cyclic, debilitating cognitive, somatic and affective symptoms (depression, irritability, mood lability, anxiety) which occur during the luteal phase, abate at menses, and greatly affect quality of life^18^. PMDD, which is now categorized as a new depressive disorder in the Diagnostic and Statistical Manual of Mental Disorders (DSM–5) (America Psychiatric Association), affects approximately 5-8% of women of reproductive age. An additional 30-40% of women suffer from milder, yet clinically significant, premenstrual symptoms (PMS), that also significantly impact the quality of life^19^. Finally, the knowledge about menstrual cycle-related emotional experiences without clinical relevance is sparse and anecdotal.

Despite the overwhelming evidence of the behavioural effects of the menstrual cycle, very little evidence is available on what the potential mechanisms might be. Classically, the connection between behavioural features and brain activity has been studied based on the assumption that specific brain areas serve specific functions. This approach successfully explains, and to some extent predicts, the impairment of relatively simple functions such as motor or sensory ones^20^. However, it has not been possible to identify specific brain locations responsible for higher cognitive function. Hence, for the understanding of higher mental functions it is necessary to adopt a more integrated approach, where the brain description is not limited to the properties of individual areas, but includes also structural and functional relationship among them, forming a tightly regulated network giving rise to the emergence of complex behaviour^21^. In the last decades, new techniques and higher available computational power made it possible to analyse brain activity non-invasively at the whole-brain level, typically through the prism of graph theory, where nodes of the graph represent brain areas, and edges represent statistical dependencies among the signals generated by such areas. These techniques can be applied to the BOLD signal derived from functional magnetic resonance (fMRI), as well as to neurophysiological (electroencephalography - EEG- and magnetoencephalography - MEG) signals.

Due to the abovementioned physiological cyclical fluctuations, both with respect to hormones levels and behaviour, the different phases of menstrual cycle can be exploited to study the relationship between sex hormones level, behavioural changes and functional brain correlates. However, the brain network research applied to the menstrual cycle highlights conflicting evidences. Using functional magnetic resonance (fMRI), Petersen et al.^22^ have demonstrated the influence of sex hormones during follicular and luteal phases on two different functional network, the anterior portion of the default mode network (aDMN) and the executive control network (ECN). In detail, comparing the brain networks in the two phases, they found that during the follicular phase, the connectivity of both the left angular gyrus within the aDMN and the right anterior cortex within the ECN were increased. Arélin et al.^23^ investigated the associations between ovarian hormones and eigenvector centrality (EC) in resting state functional magnetic resonance imaging (rs-fMRI) across menstrual cycle, and found a positive correlation between progesterone and EC in the dorsolateral prefrontal cortex, sensorimotor cortex and the hippocampus, suggesting the modulator role of progesterone on areas involved in memory regulation. In contrast, both Hjelmervik et al.^24^ and De Bond et al.^25^ did not find any changes in rs-fMRI brain connectivity in different menstrual cycle phases. The influence of sex hormones on brain connectivity was also studies through resting state electroencephalography (EEG) measurements, which provides lower spatial and higher temporal resolution as compared to fMRI. Brötzner et al.^26^ have associated the alpha frequency oscillations with menstrual cycle phases and hormones level, and found the highest alpha frequency during the luteal phase and the lowest alpha frequency during the follicular phase, with the change negatively correlated to the and estradiol levels, suggesting that the latter modulates the resting state activity in the alpha band. Finally, the correlation between brain network changes and behavioural modification typical of PMDD remains poorly explored. For example, Syan et al.^27^ examined rs-fMRI in woman with PMDD and failed to show differences in terms of brain network properties.

MEG is a non-invasive neurophysiological brain imaging method devised to measure the magnetic fields produced by the electrical activity of neuronal cells. Unlike the electric fields in the EEG, the magnetic signals are not distorted by the tissue layers surrounding the brain parenchyma, allowing high spatial accuracy^28^. Furthermore, the fact that the recorded signal is not mediated by the levels of oxygenation, as it is the case with BOLD signals, allows MEG to achieve high temporal resolution, and provide an estimate of the neuronal activity across a broad frequency spectrum. Ultimately, at present MEG is the only non-invasive brain imaging technique having at same time a sufficiently good spatial (2-5 mm) and excellent temporal (~1 msec) resolution^29^.

In this study we hypothesized that the brain network rearranges periodically along the menstrual cycle as a function of the levels of hormones. and that these changes are associated to modifications of the affective condition, even in the absence of overt clinical signs of anxiety and/or depression. To test our hypothesis, we exploited MEG to investigate the brain topology in early follicular, ovulatory and luteal phase, in twenty-four healthy, naturally-cycling women without pre-menstrual symptoms and with no signs of anxiety and/or depression. More precisely, we estimated the links between areas by invoking synchronization as a mechanism of communication^30^, and applied a novel metric, the Phase Linearity Measurement (PLM)^31^, to estimate the degree of synchronization. Once a frequency specific adjacency matrix (based on the PLM) was obtained, it was then filtered using the Minimum Spanning Tree (MST) algorithm, as to allow an unbiased comparison of topological properties between groups^32,33^. Furthermore, we correlated brain topological changes along the menstrual cycle with the corresponding changes in hormone levels (estradiol, progesterone, LH, FSH), as well as to the affective condition. The correlation between the hormonal levels and the affective condition along the menstrual cycle was investigated as well. Finally, to explore the causal relationship between hormonal levels and topological properties, we used a linear model to predict the topological properties from the hormones level.

## Results

Twenty-six women were analysed, obtaining brain topological information (computed from MEG data), blood hormone levels and psychological data at three time points across the menstrual cycle. Specifically, the participants were observed in the early follicular (T1), ovulatory (T2) and luteal (T3) phases. Two women were excluded because the BDI test value fell below the cut-off, therefore all data analysis were conducted on twenty-four women.

### Analysis of the topological parameters

In order to ascertain possible changes of brain topology across the menstrual cycle, global and nodal parameters of the brain networks (see Methods section) at early follicular, ovulatory and luteal phases have been calculated. The nodal analysis showed significant difference in the betweenness centrality (BC) of the right posterior cingulate gyrus (rPCG) (χ^2^ (df = 2, N = 24) = 15.2500, *p* = 4.8 × 10-4, *p*_FDR_ (False Discovery Rate – see methods section) = 0.043), in the alpha band. In detail, the post-hoc analysis showed significantly higher BC in the rPCG during the ovulatory phase, as compared to the follicular (*p* = 0.0003) and luteal (*p* = 0.0055) phases (Fig. 1).

**Fig. 1.**
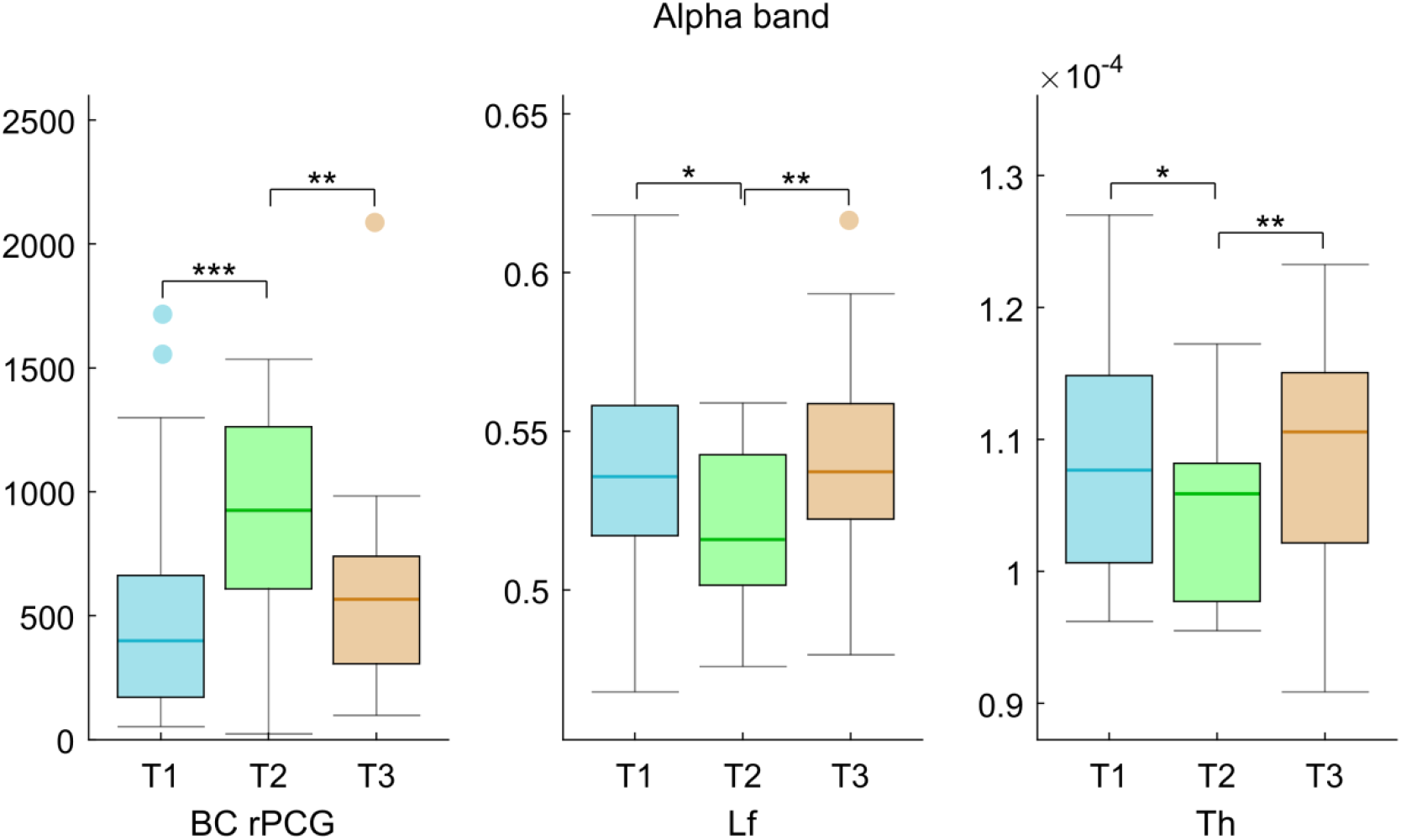
Brain topology comparison. The box plots refer to the alpha band, from left to right, to the BC in the rPCG, the leaf fraction (Lf) and the Tree hierarchy (Th), respectively. In each box plot, the values are shown at early follicular (T1), ovulatory (T2) and luteal (T3) phases. The upper and lower bound of the rectangles refer to the 25^th^ to 75^th^ percentiles, the median value is represented by horizontal line inside each box, the whiskers extent to the 10^th^ and 90^th^ percentiles, and further data are considered as outliers and represented by the filled circles. Significance *p* values: \**p* < 0.05, \*\**p* < 0.01, \*\*\**p* < 0.001.

With regard to the global topological parameters, the leaf fraction (Lf, a measure of network integration-see discussion) (χ^2^ (df = 2, N = 24) = 10.7500, *p* = 0.0046, *p*_FDR_ = 0.009) and the tree hierarchy (Th, a measure of the trade-off between a well-connected network that is also resilient to targeted attacks - see discussion) (χ^2^ (df = 2, N = 24) = 12.3333, *p* = 0.0021, *p*_FDR_ = 0.008) were reduced in the alpha band. More specifically, the post-hoc analysis revealed a reduction in the network integration in ovulatory phase as compared to both the follicular (Lf *p* = 0.016; Th *p* = 0.032) and the luteal (Lf *p* = 0.004; Th *p* = 0.006) phases. No statistically significant difference, after FDR correction, was found in any other nodal and global parameter, nor in any other frequency band. The topological parameters, that showed significant variations during the menstrual cycle (BC in the rPCG, Lf and Th) became parameter of interest for follow-up correlation and linear model analyses.

### Topological brain network parameters and hormone blood levels

To explore the possible influence of sex hormones on the topological brain configuration throughout the menstrual cycle, Spearman’s correlation analyses between the variations (Δ T1-T2 and Δ T2-T3) of the brain network topological parameters and the concurrent variations in the hormonal levels have been studied (Fig. 2). A statistically significant direct correlation between the Δ values of the BC of the rPCG in the alpha band, and those of estradiol (r = 0.67, *p* =3.6 × 10^−7^, *p*_FDR_ = 1.4 × 10^−6^), LH (r = 0.50, *p* = 0.0003, *p*_FDR_ = 0.0006) and FSH (r = 0.43, *p* = 0.0023, *p*_FDR_ = 0.0031). No correlation was demonstrated between the global topological parameters and hormonal levels.

**Fig. 2.**
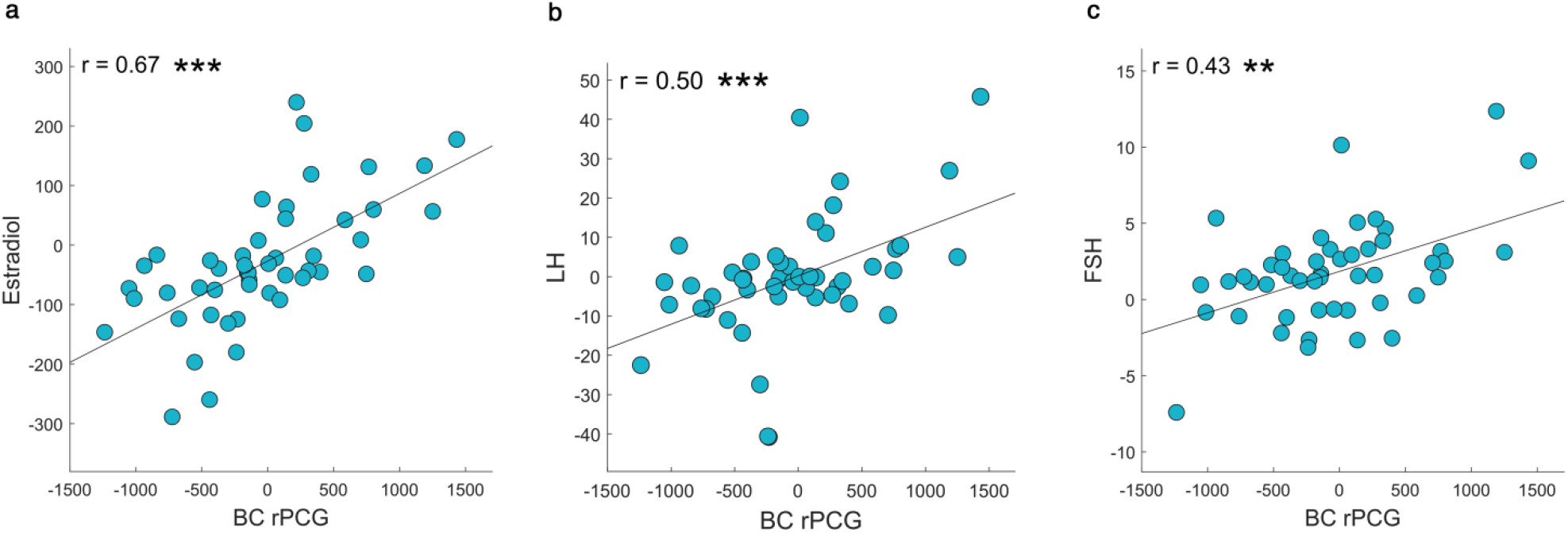
Correlation between topological data and hormones blood levels. Spearman’s correlation between the Δ values (expressed by the variation between the menstrual cycle phases (Δ t1-t2 and Δ T2-T3)) of betweenness centrality (BC) of the right posterior cingulate gyrus (PCG) and the Δ values of (**a**) estradiol, (**b**) luteinizing hormone (LH), and (**c**) follicular stimulant hormone (FSH) levels along the menstrual cycle. Significance *p* values: \**p* < 0.05, \*\**p* < 0.01, \*\*\**p* < 0.001.

### Topological brain network parameters and psychological scores

To study whether the topological changes that we have observed could be linked to the well documented changes of the affective condition across the menstrual cycle, Spearman’s correlation analysis between the brain network parameters and the psychological scores were carried out (Fig. 3). The analysis showed a significant direct correlation between the Δ values of the BC of the rPCG and the Δ values of two of the six subdomains of the well-being test, namely the environmental mastery (*r* = 0.40, *p* = 0.004, *p*_FDR_ = 0.013) and the self-acceptance (*r* = 0.42, *p* = 0.002, *p*_FDR_ = 0.013). No correlation was demonstrated with the global topological parameters.

**Fig. 3.**
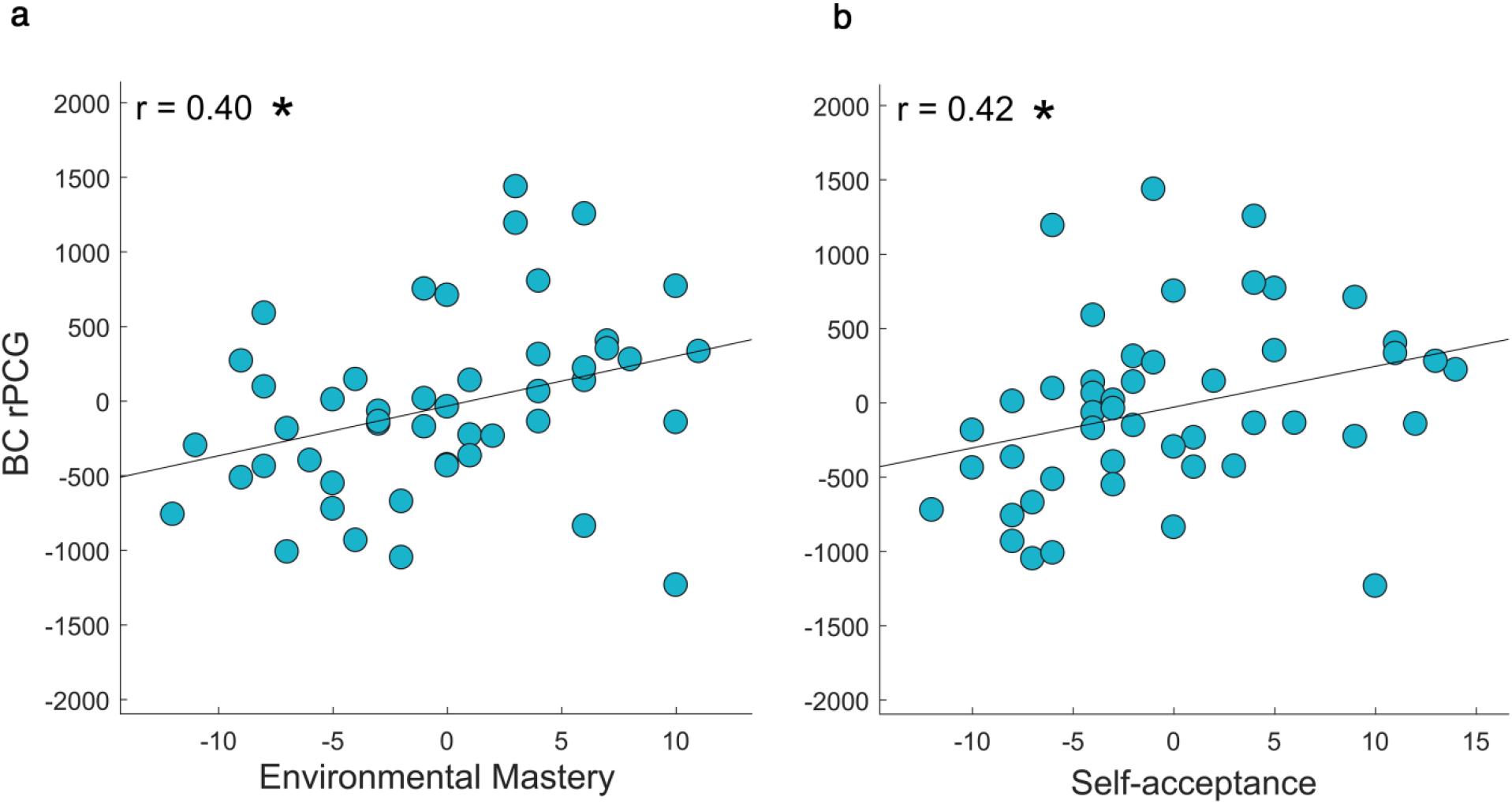
Correlation between topological data and psychological dimensions of Well-Being test. Spearman’s correlation between the Δ values (expressed by the variation between the menstrual cycle phases (Δ T1-T2 and Δ T2-T3)) of betweenness centrality (BC) of the right posterior cingulate gyrus (PCG) and the Δ values of psychological dimensions of the Well-Being test ((**a**) Environmental Mastery and (**b**) and Self-acceptance scores) along the menstrual cycle. Significance *p* values: \**p* < 0.05, \*\**p* < 0.01, \*\*\**p* < 0.001.

### Hormone blood levels and psychological scores

To analyse the possible association between sex hormones changes and the well-known affective modifications occurring during the menstrual cycle, Spearman’s correlation analysis between the Δ values of the hormonal levels and those of the psychological scores was performed (Fig. 4). A statistically significant correlation between estradiol and environmental mastery (*r* = 0.44, *p* = 0.001, *p*_FDR_ = 0.034) was observed.

**Fig. 4.**
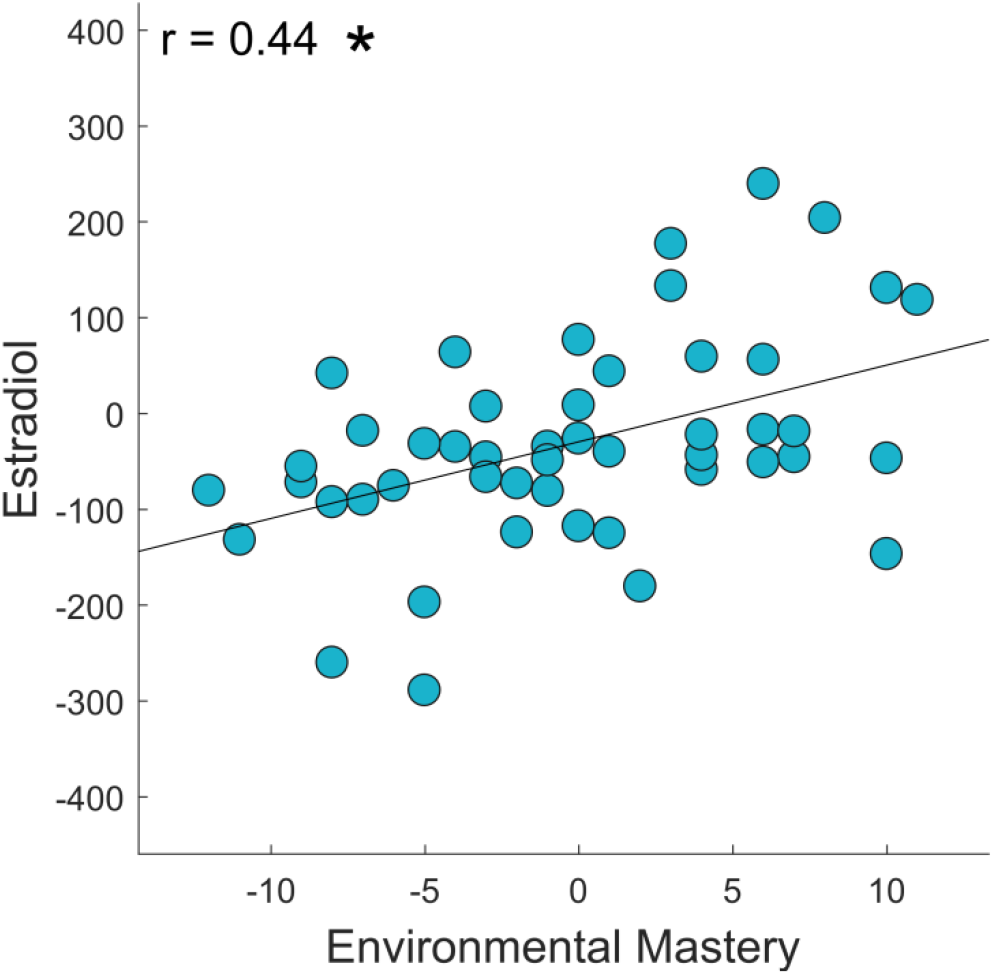
Correlation between hormones blood levels and psychological dimensions of Well-Being test. Spearman’s correlation between the Δ values (expressed by the variation between the menstrual cycle phases (Δ T1-T2 and Δ T2-T3)) of Estradiol and the Δ values of psychological dimension of Well-Being test (Environmental Mastery scores) along the menstrual cycle. Significance *p* value: \**p* < 0.05, \*\**p* < 0.01, \*\*\**p* < 0.001.

### Multilinear model analysis

To further explore the causal relationship occurring between changes of topological properties in the brain and hormonal levels, we build a linear model to predict the changes (Δ T1-T2 and Δ T2-T3) in the BC in the rPCG, and in the Lf and the Th, as a function of the changes in the hormonal levels, using the leave-one-out cross-validation approach (LOOCV) (see Methods for details) (Fig. 5). Hormone blood level variations (Δ T1-T2 and Δ T2-T3) of estradiol, progesterone, LH, FSH scores were included into an additive multilinear model, together with nuisance variables (age, education, cycle length). We found that the model yielded significant predictions of the BC of the rPCG (R^2^ = 0.51), with estradiol being a significant predictor for the model (p < 0.001), with positive beta coefficients. The prediction of the model and the distribution of the residuals (computed through LOOCV) are shown in Fig. 5, panels B and C. The same model was applied to global topological parameters, but no significant results were obtained.

**Fig. 5.**
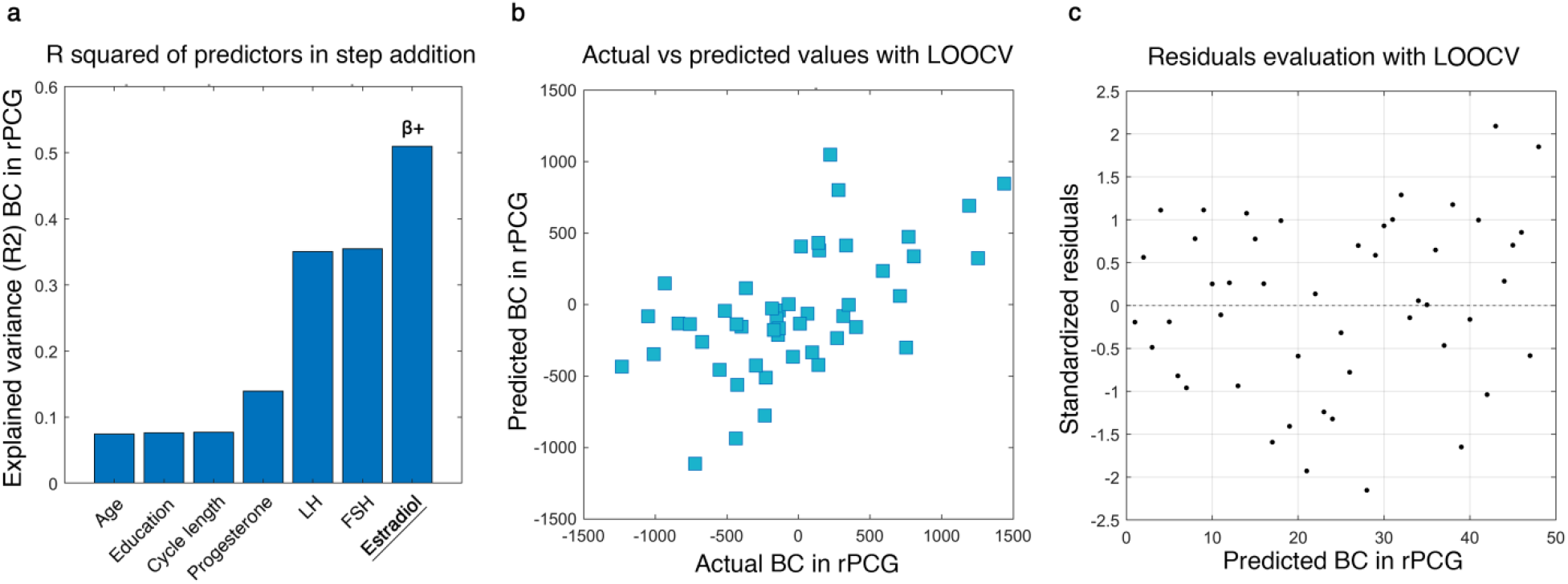
Multilinear model with leave-one-out cross-validation (LOOCV). The model aims to predict the topological variation of the brain network expressed by the betweenness centrality (BC) changes during the menstrual cycle (Δ T1-T2 and Δ T2-T3) of the right posterior cingulate gyrus (rPCG). **a**, Explained variance of the additive model composed of three nuisance variables (age, education, cycle length), and four predictors (progesterone, luteinizing hormone (LH), follicle-stimulating hormone (FSH), estradiol). Significant predictor in underlined text; positive coefficient indicated with β+. **b**, Scatter plot of the Observed topological values versus the topological values predicted by the model with LOOCV. **c**, Scatter plot of the standardized residuals (standardization of the difference between observed and predicted (LOOCV) values). The distribution results symmetrical with respect to the 0, with a standard deviation lower than 2.5.

## Discussion

In the present study, we set out to test the hypothesis that sex hormones changes, as they occur across the menstrual cycle, may affect the topological configuration of brain networks, as well as modulate the frequently observed mood changes. We showed that during the menstrual cycle the topological features of the brain network undergo profound rearrangements under the effect of sex hormones, as highlighted by changes in both nodal and global topological parameters. In particular, we showed in the alpha band, during the ovulatory phase, increased BC in the right posterior cingulate gyrus and reduced Lf and Th, as compared to both the follicular and luteal phases. The increase of the BC of the right posterior cingulate gyrus was positively correlated with the changes in the blood levels of estradiol, LH and FSH. Furthermore, though a multilinear model, we showed that nearly 50% of the variance of the changes of the BC can be explained by the estradiol. We also demonstrated that the increase in the BC was related to improved subjective well-being, as suggested by the positive correlation to the scores of the environmental mastery and the self-acceptance domains within the well-being test. Finally, we showed that the environmental mastery domain of the well-being test was also correlated to the estradiol levels.

The PCG is described as “an enigmatic cortical region”^34^. If, on the one hand, the high metabolic expenditure and the number of cortical and subcortical connections point at the PCG as structural and functional hub, on the other hand, growing evidence shows that the PCG tends to deactivate in response to attention demanding tasks^35^. Accordingly, the PCG displays increased activity when the subject is involved in internally-directed task such as retrieving autobiographical memories, planning for the future or wandering freely with the mind^36–38^. Recent studies suggest that the PCG may play a crucial role in the stepwise mechanisms of integration of specialized perceptive processes (i.e. visual, auditory or sensory) into higher levels of abstraction. Other works suggest that the PCG may play a role in assessing the significance of decision outcome, being important in balancing between risk-prone and risk-adverse behaviours^34^.

It is noteworthy that the PCG change is not symmetric. This fact may be associated with a different influence of the sex hormones on the right and left PCG, possibly modulating the expression of affective behavioural styles. Hwang et al.^39^ demonstrated an asymmetry on the way the brain is modulated by sex hormones during the menstrual cycle. In particular, higher right frontal activity was observed during the ovulation phase, and a higher left activity during the menstruation phase. However, they found this left-right asymmetry in the frontal regions of the brain, while our data points at the posterior brain regions. Nonetheless, it is interesting to note that the DMN areas possess long-distance projections to the anterior cingulate areas via the PCG^40^. Furthermore, has been shown that the asymmetry at rest between the right and left sides of brain represents a reliable measure of individual affective style^41^. In particular, greater alpha activity in the right regions corresponds to a personality trait sensitive to negative affective stimuli, while greater alpha activity on the left corresponds to a personality trait sensitive to positive affective stimuli.

Furthermore, we showed a statistical significant reduction of the Lf and the Th during the ovulatory phase, as compared to the follicular and luteal phases. This data might suggest a shift towards a less centralized organization of the brain network^42^ in which the information flow is less reliant on any single node, with consequent improved resiliency to targeted attacks^32,33^. These results could be summarized as a better global efficiency which is an expression of an optimal organization of the brain network during the ovulatory phase, in terms of an optimal trade-off between efficient communication and resiliency.

The direct correlation between the BC changes in the rPCG and the variations of estradiol, LH and FSH suggests that the monthly hormonal fluctuations affect the role of this area within the brain networks. Our results suggest that the sex hormones changes, and specifically those involving estradiol, LH and FSH, have a substantial impact on the functional architecture of the brain networks. Besides the evidence of higher BC in the rPCG during the ovulation phase, we also show that the changes of BC are linearly proportional to the changes of the blood estradiol level. This fits with the fact that estradiol, LH and FSH levels peak during the ovulatory phase.

The multilinear model confirmed that there is a relationship between the topological variation and the hormonal fluctuations that occur during the menstrual cycle, in fact nearly 50% of the variance of the changes of the BC in the PCG during the menstrual cycle can be explained by the changes in estradiol. Multiple works have tried to disentangle hormone-specific influences on the brain networks. However, the literature is largely inconsistent, even when limiting oneself to the effects of hormones on brain connectivity alone. Several studies have shown the involvement of the estradiol on both the structure and the function of the brain. In particular, it has been observed that estradiol affects the activity of the right anterior hemisphere^39^. Pletzer et al.^43^ demonstrated that the left hippocampus is highly activated during the pre-ovulatory phase, while its activation drops during the luteal phase, suggesting that estradiol and progesterone have opposite effects on the hippocampus. Furthermore, MRI studies have reported increased grey matter volumes in the hippocampus during the pre-ovulatory phases^43,44^. A resting state MRI study found a significant positive correlation between progesterone (but not estradiol) and the Eigenvector centrality in the dorsolateral prefrontal cortex in a single woman scanned 32 times across four menstrual cycles^23^. However, further studies did not find any correlation between resting state activity and neither progesterone nor estradiol^22^. Very recently, Pritschet et al.^45^ demonstrated, in a very elegant study, the crucial effect of estradiol on brain network. The authors performed a dense-sampling protocol, scanning the same woman for 30 consecutive days. One year later the same woman repeated the protocol while she was under hormonal therapy, as to selectively suppress progesterone synthesis, while leaving estradiol unaffected. In the second experimental setting, the authors were able to confirm the previous results.

Our observation about the positive correlation between the increase in BC, suggesting a greater topological centrality of the rPCG within the cerebral network, and higher levels of estrogen, LH and FSH, does not find an immediate and unambiguous explanation. Albeit within a purely speculative framework, we notice that the greater centrality of the PCG is coupled to the levels of estradiol, LH and FSH, showing that the role of this region within the network is more prominent during the moment of fertility. Observing this phenomenon from an evolutionistic perspective, one could think that, when fertility is at its peak a quick and effective evaluation of the relative risks and rewards associated to the potential mate would be adaptive. The PCG might have implications in the top-down control in decision making as in the choice of the partner^46,47^.

Several studies support this hypothesis. For example, an event-related potential, source reconstructed EEG study^48^ reveals that the strongest activations were in the PCG when presenting scenes in which 2 people performed “affective” actions, while the superior temporal sulcus, an area included in the mirror neuron system, was activated by cooperative scenes. In fact, it is well established that the PCG is involved in emotion processing^49,50^, in the subjective evaluation of events, and in the attribution of their emotional significance. Furthermore, this observation seems to be gender-specific, since women show improved comprehension of unattended social scenes as compared to men. Rupp et al.^51^ used fMRI to measure brain activity in twelve women as they evaluated pictures of masculinized or feminized male faces, during both the follicular and luteal phase. They found that the brain regions involved in face perception, decision making and reward processing, including the PCG, responded more strongly to masculinized faces as compared to feminized ones. Additionally, the authors showed that such process was influenced by the hormonal levels. More specifically, the PCG activation was positively predicted by estradiol (and testosterone). They propose that this mechanism may have a role in the women’s cognitive processes underlying the decision making process in partner choice. Further extensive literature is available to sustain this hypothesis^52–59^.

An experience shared by a very large number of women of childbearing age is an emotional lability during the luteal phase, in the days immediately before the menstruation^60^. This condition can take on clinical relevance in the form of PMS or even grow to a dysphoric clinical picture as in the case of PMDD^16,17^. A number of studies have investigated the role of sex hormones in PMS/PMDD, but no abnormal levels have been established^61,62^ although with inconsistency^63^. At moment, the hypothesis with the stronger consensus claims a maladaptive response of the brain regions involved in affective processes to the physiological fluctuations of the sex hormones^64^. PMS/PMDD would result from an imbalance between bottom-up processes, involving the amygdala and the insular cortex, and top-down regulation through the prefrontal and cingulate cortices. In this study, we sought to provide evidence about the possible correlation between clinically under-threshold affective modifications and both topological changes and sex hormone fluctuations observed along the menstrual cycle. We showed a positive association between the BC values of rPCG and the subjective well-being in the environmental mastery and self-acceptance sub-domains of the Ryff’s test. Furthermore, we showed a correlation between the environmental mastery sub-domain of the Ryff’s test and estradiol levels. Our data demonstrate that during the ovulatory phase, when estradiol reaches its peak, the BC values of the rPCG peak as well. At same time, the affective state correlates positively with both the BC of the rPCG and estradiol blood levels. The combination of these observations (the positive correlation between the BC of the rPCG and both sex hormones and affective state, and the correlation between estradiol and affective condition) suggests that the sex hormones interfere with the affectivity, possibly by changing the topological features of the rPCG, a brain region specifically involved in the top-down computation of emotional stimuli^34^.

In conclusion, the responsiveness to affective and emotional stimuli is not constant. Rather, it may be accentuated or attenuated during the menstrual cycle perhaps via the modulation of the sex hormones, trough mechanisms acting at different levels, including rearrangements of the large-scale functional architecture of the brain. The results we present provide relevant information for all the studies that use brain topological indices to compare multiple groups. In fact, provided that the topological parameters are influenced by the hormonal profile, at least in women, this information should be taken into consideration to avoid a biased comparison.

## Conclusions

In conclusion, we have shown that during the ovulatory phase an increase in the values of BC in the rPCG occurs. The changes in BC correlate positively with the estradiol, LH and FSH blood levels, all of which have their concentration peak in the ovulatory phase. The multilinear regression analysis confirmed that there is a strong relationship between the topological variation and estradiol. We have also highlighted how high BC values in the rPCG are linked to a better affective condition as suggested by the positive correlation with tests that evaluate the well-being in the dimensions of environmental mastery (and self-acceptance) that, in turn, is correlated to the estradiol levels. Finally, our work has widespread implications for all clinical neuroimaging studies, given that the comparison between groups should account for the physiological variations in the brain topology that occur in women throughout the menstrual cycle

## Methods

### Participants

Twenty-six strictly right-handed, native Italian speaker females were recruited (Tab. 1). We included women with a regular menstrual cycle (mean cycle length 28.4 ± 1.3 days), who had not make use of hormonal contraceptives (or other hormone regulating medicaments) during the last six months before the recording, who had not been pregnant in the last year and, finally, without history of neuropsychiatric diseases or premenstrual dysphoric/depressive symptoms. To check for mood and/or anxiety symptoms, the Beck Depression Inventory (BDI)^65^ and Beck Anxiety Inventory (BAI)^66^ were used with a cut-off below 10 and 21, respectively. To control for influence of circadian rhythm, the time of testing varied no more than two hours between testing sessions. To control for a possible session effect, women were randomized according to the cycle phase at the first session.

**Tab. 1.** Demographic and anamnestic data. Data are given as mean ± standard deviation (SD).

| Demographic and anamnestic data |  |
| --- | --- |
| *Parameters* | *Participants (24)* |
| Age (years) | 26.2 ± 5.1 |
| Education (years) | 17.1 ± 2.7 |
| Menstrual cycle duration (days) | 28.4 ± 1.3 |

### Experimental protocol

At enrolment, all women signed a written consent form. All the procedures strictly adhered to the guidelines outlined in the Declaration of Helsinki, IV edition. The study protocol was approved by the local ethic committee (University of Naples Federico II; protocol n. 223/20). Demographic and anamnestic data were collected and recorded on a dedicated database (Tab. 1). The women were tested in three different cycle phases, i.e. in the early follicular phase (cycle day 1-4, low estradiol and progesterone, T1), during the ovulatory phase (cycle day 13-15, high estradiol, T2) and in luteal phase (cycle day 21-23, high estradiol and progesterone, T3). To estimate individual cycle phases, the back-counting method was applied. Self-reported onset of menses was used as a starting point. During the three times cycle, all subjects underwent the following examinations: MEG recording, blood sampling for the hormone dosage and psychological evaluation. During the follicular phase a transvaginal pelvic ultrasonography examination was performed. After the last MEG recording, a structural magnetic resonance imaging (MRI) was performed. Two subjects refused to execute the MRI scan and consequently the template was used for sources reconstruction.

### Ultrasound examination

All participants underwent a transvaginal pelvic ultrasonography during the early follicular phase. Scans were performed using a 4-10 MHz endocavitary transducer (GE Healthcare, Milwaukee, WI). Patients were in lithotomy position with empty bladder. The uterus and both ovaries were visualized. The uterus was scanned using longitudinal and transverse plane, endometrial thickness was measured at the widest point in the longitudinal plane. Follicle number and diameters were assessed for each ovary. Presence of abnormal findings, such as endometrial polyps, myomas, ovarian cysts or other adnexal masses was addressed. None of the enrolled patients presented abnormal findings and endometrial thickness and follicle diameters were consistent with the menstrual phase.

### Hormone assays

Each participant underwent venous blood sampling during the three hormonal phases of the menstrual cycle. All women were asked to respect a 12-h fast before blood collection. Whole blood samples were collected in S-Monovette tubes (Sarstedt), containing gel with clotting activator in order to facilitate the separation of the serum from the cellular fraction, according to predetermined standard operating procedure^67^. To this aim, samples were centrifuged at 4000 rpm for 10 minutes, then the serum was collected, aliquoted in 1.5 ml tubes (Sarstedt) and stored at –80 °C until the analysis. Determination of estradiol (range: 19,5-144,2 pg/ml (follicular phase); 63,9-356,7 pg/ml (ovulatory phase); 55,8-214,2 pg/ml (luteal phase); detection limit: 11,8 pg/mL; inter-assay coefficients of variation averaged: 1,9%; Intra-assay coefficients of variation averaged: 4,9%), progesterone (range: ND-1,4 ng/ml (follicular phase); ND-2,5 ng/ml (ovulatory phase); 2,5-28,03 ng/ml (luteal phase); detection limit: 0,2 ng/ml; inter-assay coefficients of variation averaged: 5,5%; intra-assay coefficients of variation averaged: 3,56%), LH (range: 1,9-12,5 mIU/ml (follicular phase); 8,7-76,3 mIU/ml (ovulatory phase); 0,5-16,9 mIU/ml (luteal phase); detection limit: 0,07 mIU/ml; inter-assay coefficients of variation averaged: 2,3%; intra-assay coefficients of variation averaged: 2,5%) and FSH levels (range: 2,5-10,2 mIU/ml (follicular phase); 3,4-33,4 mIU/ml (ovulatory phase); 1,5-9,1 mIU/ml (luteal phase); detection limit: 0,3 mIU/ml; inter-assay coefficients of variation averaged: 1,2%; intra-assay coefficients of variation averaged: 1,9%) were measured by Advia Centaur XT Immunoassay System analyzer (Siemens) which uses competitive (estradiol) or direct (progesterone, FSH, LH) immunoassay and for quantification of reaction uses Chemiluminescent Acridinium Ester technology. The 2.5^th^ and 97.5^th^ percentiles were used to form reference limits with 90% confidence intervals, as provided by assay manufacturers^68^.

### MEG recording

The MEG system was developed by the National Research Council (CNR), Pozzuoli, Naples, at Institute of Applied Sciences and Intelligent Systems “E. Caianiello”, and is placed inside a shielded room (AtB Biomag UG-Ulm–Germany). The MEG is equipped by 154 magnetometers and 9 reference sensors located on a helmet^69^. Before each MEG session, four position coils were placed on the participant’s head and their position, as well as that of four anatomical landmarks, was digitized using Fastrak (Polhemus®)^70^. The coils were activated and localized at the beginning of each segment of registration. The magnetic fields were recorded for 7 minutes, divided into two time intervals of 3’30’’, while the participants were sitting comfortably in an armchair in the cabin with their eyes closed, and were instructed not to think of something in particular. During the acquisition the electrocardiogram (ECG) and electro-oculogram (EOG) signals were also recorded^71^. The data were sampled at *fs* = 1024 Hz and a 4^th^ order Butterworth IIR pass-band filter between 0.5 and 48 Hz was applied. After each session, all the subjects were checked for drowsiness during the recording with a specific questionnaire.

### Data processing and source reconstruction

After the recording phase, the brain magnetic signals were cleaned through an automated process as described in our previous article^72^. The FieldTrip software tool^73^, based on Mathworks® MATLAB, was used to implement principal component analysis (PCA)^74,75^, to reduce the environmental noise, and independent component analysis (ICA)^76^, to remove physiological artefacts such as cardiac noise or eyes blinking (if present). For each participant, source reconstruction was performed for all segments through a beamforming procedure using the Fieldtrip toolbox similarly to Jacini et al.^77^. In short, based on the native MRI, the volume conduction model proposed by Nolte^78^ was applied and the Linearity Constrained Minimum Variance beamformer^79^ was implemented to reconstruct time series related to the centroids of 116 regions-of-interest (ROIs), derived from the Automated Anatomical Labeling (AAL) atlas^80,81^. We only considered the first 90 ROIs, excluding those corresponding to cerebellum, given that the reconstructed signal might be less reliable.

### Construction of brain network

After the signal had been filtered in each canonical frequency band (i.e. delta, theta, alpha, beta and gamma – see later), the Phase Linearity Measurement (PLM)^82^ was computed, to provide an estimate of synchronization between any two region that is purely based upon the phases of the signals, and unaffected by volume conduction. The PLM is defined as^31^:

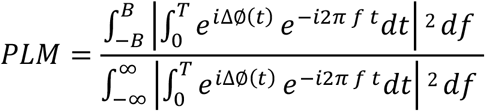

where the Δ∅(*t*) represent the phase difference between two signals, the 2B is the frequency band range, set to 1 Hz, *f* is the frequency and *T* is the observation time interval.

The PLM was performed for segments longer than 4s. By computing the PLM for each couple of brain regions, we obtained a 90×90 weighted adjacency matrix for each time series and for each subject, in all frequency bands: delta (0.5–4 Hz), theta (4.0–8.0 Hz), alpha (8.0–13.0 Hz), beta (13.0–30.0 Hz) and gamma (30.0–48.0 Hz). Each weighted adjacency matrix was used to reconstruct a brain network^32^, where the 90 areas of the AAL atlas are represented as nodes, and the PLM values form the weighted edges. For each trials longer than 4s, and for each frequency band, through Kruskal’s algorithm^83^, the minimum spanning tree (MST) was calculated. The MST is a loop-less graph with N nodes and M = N-1 links. The MST was computed to be able to compare topological properties in an unbiased manned^32,33^.

### Graph analysis

Global and nodal (regional) parameters were calculated. In order to characterize the global topological organization of the brain networks, four topological parameters were calculated. The *leaf fraction* (Lf)^42^, defined as the fraction of nodes with a degree of 1, provides an indication of the integration of the network, with high leaf fraction conveying a more integrated network. The *degree divergence* (K)^42^, a measure of the broadness of the degree distribution, is related to the resilience against targeted attacks. The *tree hierarchy* (Th)^42^ is defined as the number of leaf over the maximum betweenness centrality, and is meant to capture the optimal trade-off between network integration and resiliency to hub failure. Finally, the *diameter*^42^ is defined as the longest shortest path of an MST, and represent a measure of ease of communication flow across a network. To examine the relative importance of specific brain areas in the brain network, two centrality parameters were calculated: the *degree*^33^, defined as the number of edges incident on a given node, and the *betweenness centrally* (BC)^33^, defined as the number of the shortest paths passing through a given node over the total of the shortest paths of the network. Before moving to the statistical analysis, all the metrics were averaged across epochs to obtain one value for subject. A pipeline of the processing MEG data is illustrated in Fig. 6.

**Fig. 6.**
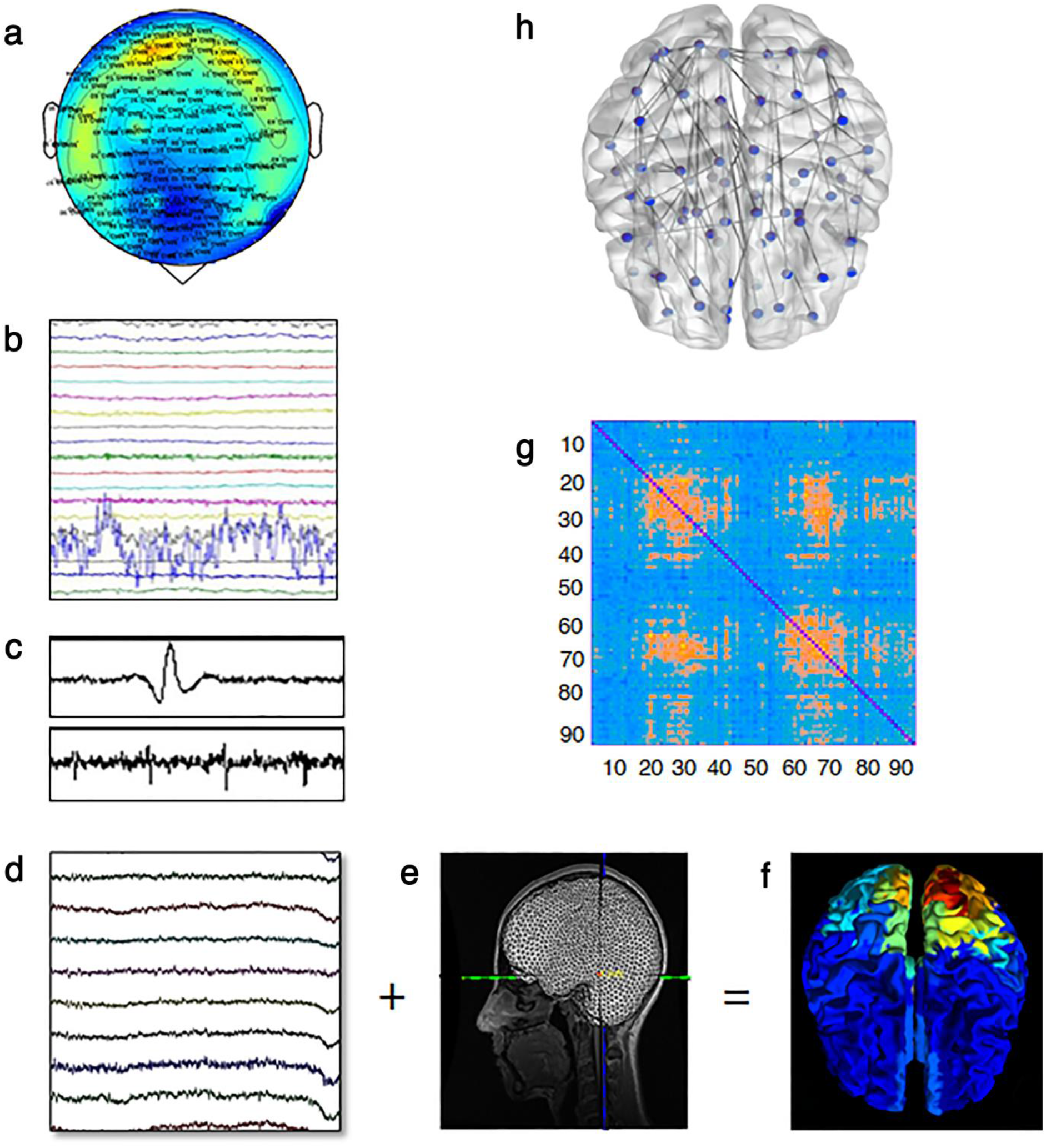
Data analysis pipeline. **a**, Neuronal activity recorded by 163 sensors. **b,** Noisy channels identified by an experienced rater. **c**, Cardiac (upper) and blinking (lower) artefacts as estimated by ICA. **d**, Cleaned channels. **e**, Native MRI. **f**, Structural MRI and MEG sensors are co-registered and the time series are estimated in source space. **g**, Functional connectivity matrix estimated using the Phase Linearity Measurement. Rows and columns are the regions of interest, while the entries are the estimated values of the Phase Linearity Measurement. **h,** Brain topology representation based on the MST.

### MRI acquisition

MRI images of twenty-four participants were acquired on a 1.5-T Signa Explorer scanner equipped with an 8-channel parallel head coil (General Electric Healthcare, Milwaukee, WI, USA). In particular, three-dimensional T1-weighted images (gradient-echo sequence Inversion Recovery prepared Fast Spoiled Gradient Recalled-echo, time repetition = 8.216 ms, TI = 450 ms, TE = 3.08 ms, flip angle = 12, voxel size = 1×1×1.2 mm1; matrix = 256×256) were acquired. Two subjects refused to perform MRI scan and a standard template was used to sources reconstruction.

### Psychological evaluation

The psychological assessments were carried out at each of the three menstrual cycle phases. In particular to quantify the self-esteem level, the Rosenberg Self-Esteem Scale^84,85^ was used. Additionally the Ryff’s test^86^ was administrated to examine the psychological well-being of all participants. Finally, in addition to BAI^66^ and BDI^65^ tests administrated at the first experimental session (as inclusion/exclusion criteria), the tests were re-administrated at each time point to exclude the appearance of depressive/anxious symptoms. Two women were excluded because the BDI test value had dropped below the cut-off.

### Statistical analysis

Statistical analysis was performed using MATLAB (Mathworks®, version R2013a). The normal distribution of variables was checked through the Shapiro-Wilk test. In order to compare, in all frequency bands, the topological data among the three phases of the menstrual cycle, we used the Friedman test. All the *p* values were corrected for multiple comparison using the false discovery rate across parameters for each frequency bands (FDR)^87^. Subsequently, the post-hoc analysis was carried out using Wilcoxon test. The statistical significance was defined as *p* < 0.05.

If a topological parameter was statistically different in a time point of the menstrual cycle (as compared to the other time points), we went on to check if its variation across the time points were proportional to the hormonal variations. To do this, we calculated the delta values (Δ), expressed by the variations between the menstrual cycle phases (Δ T1-T2 and Δ T2-T3) for the topological parameters (namely, the BC in the right posterior cingulate gyrus, the Lf and the Th), and the hormonal variations (estradiol, FSH, LH, and progesterone) across the same time-points. Finally, the changes of the scores of the psychological tests (self-esteem and well-being with the six relative subdomains) were related to the topological changes, as well as to the hormonal variations. The correlation analysis was performed through the Spearman’s correlation test, and the *p* values were corrected for multiple comparisons using FDR across metrics and frequency bands. A (corrected) *p* value < 0.05 was accepted as significant.

To test the hypothesis that, during the menstrual cycle, the hormonal changes provoke the topological changes, we build a linear model to predict the topological values based on hormones. Specifically, we considered the topological variation (Δ T1-T2 and Δ T2-T3) as the dependent variable, while estradiol, progesterone, LH, FSH variation (Δ T1-T2 and Δ T2-T3) were set as predictors. Moreover, in order to take into account for the possible effects of age, education and menstrual cycle length, we added these three nuisance variables as predictors too. To make the prediction of our model more reliable and to test its generalization capacity, we used a leave-one-out cross-validation (LOOCV) technique. Expressly, we built *n* multilinear model (where *n* is the size of the sample included in the model), excluding each time a different element from the model, and verifying the ability of the model to predict the topological value of the excluded element.

## Data Availability

The data used to support the findings of this study are available from the corresponding author upon request.

## Acknowledgments

The present research was supported by the University of Naples Parthenope “Ricerca locale” (GS).

## Author contributions

M. L. collected the sample, performed the MEG recordings, analysed the data, wrote the manuscript and prepared the figures. E. TL. performed the MEG recordings, designed the multilinear model, analysed the data and wrote the manuscript. L. S. performed the ultrasound examination. R. R. performed the MEG recordings and contributed to data analysis. R. M. performed the MEG recordings. M. P. collected the psychological data. G. P. and P. F. performed the pre-processing of hormones assay. F. L. provided critical revisions of manuscript. G. S. collected the venous blood sampling, contributed to interpreting the results and wrote the manuscript. P. S. supervised the study, designed the multilinear model, contributed to interpreting the results and wrote the manuscript.

